# PhenomeXcan: Mapping the genome to the phenome through the transcriptome

**DOI:** 10.1101/833210

**Authors:** Milton Pividori, Padma S. Rajagopal, Alvaro Barbeira, Yanyu Liang, Owen Melia, Lisa Bastarache, YoSon Park, The GTEx Consortium, Xiaoquan Wen, Hae K. Im

**Author notes:** Both authors contributed equally to this manuscript (Milton Pividori, Padma S. Rajagopal). Please see Supplementary Materials for full author list (The GTEx Consortium).

## Abstract

Large-scale genomic and transcriptomic initiatives offer unprecedented ability to study the biology of complex traits and identify target genes for precision prevention or therapy. Translation to clinical contexts, however, has been slow and challenging due to lack of biological context for identified variant-level associations. Moreover, many translational researchers lack the computational or analytic infrastructures required to fully use these resources. We integrate genome-wide association study (GWAS) summary statistics from multiple publicly available sources and data from Genotype-Tissue Expression (GTEx) v8 using PrediXcan and provide a user-friendly platform for translational researchers based on state-of-the-art algorithms. We develop a novel Bayesian colocalization method, fastENLOC, to prioritize the most likely causal gene-trait associations. Our resource, PhenomeXcan, synthesizes 8.87 million variants from GWAS on 4,091 traits with transcriptome regulation data from 49 tissues in GTEx v8 into an innovative, gene-based resource including 22,255 genes. Across the entire genome/phenome space, we find 65,603 significant associations (Bonferroni-corrected p-value of 5.5 × 10^−10^), where 19,579 (29.8 percent) were colocalized (locus regional colocalization probability > 0.1). We successfully replicate associations from PheWAS Catalog (AUC=0.61) and OMIM (AUC=0.64). We provide examples of (a) finding novel and underreported genome-to-phenome associations, (b) exploring complex gene-trait clusters within PhenomeXcan, (c) studying phenome-to-phenome relationships between common and rare diseases via further integration of PhenomeXcan with ClinVar, and (d) evaluating potential therapeutic targets. PhenomeXcan (phenomexcan.org) broadens access to complex genomic and transcriptomic data and empowers translational researchers.

**One-Sentence Summary:** PhenomeXcan is a gene-based resource of gene-trait associations with biological context that supports translational research.

## Introduction

Unprecedented advances in genetic technologies over the past decade have identified over tens of thousands of variants associated with complex traits (*1*). Translating these variants into actionable targets for precision medicine or drug development, however, remains slow and difficult *(2)*. Existing catalogs largely organize associations between genetic variants and complex traits at the variant level rather than by genes, and often are confined to a narrow set of genes or traits (*3*). This has greatly limited development and application of large-scale assessments that account for spurious associations between variants and traits. As a result, only 10 percent of genes are under active translational research, with a strong bias towards monogenic traits (*4,5*).

Complex diseases are generally polygenic, with many genes contributing to their variation. Concurrently, many genes are pleiotropic, affecting multiple independent traits *(6)*. Phenome-wide association studies (PheWAS) aim to complement genome-wide association studies (GWAS) by studying pleiotropic effects of a genetic variant on a broad range of traits. Many PheWAS databases aggregate individual associations between a genetic variant and a trait, including GeneATLAS (778 traits from the UK Biobank (http://geneatlas.roslin.ed.ac.uk/trait/)) *(7)*, GWAS Atlas (4,155 GWAS examined over 2,965 traits (https://atlas.ctglab.nl/)) (*8*), and PhenoScanner (over 5,000 datasets examined over 100 traits (http://www.phenoscanner.medschl.cam.ac.uk/)) (*9*). Other PheWAS databases are constructed based on polygenic scores estimated from multiple variants per GWAS locus (*10)*, latent factors underlying groups of variants (*11*) or variants overlapping between GWAS and PheWAS catalogs (*12*). By building associations directly from variants (most of which are non-coding), most PheWAS results lack mechanistic insight that can support proposals for translational experiments. Genes are primarily assigned to PheWAS results by genomic proximity to significant variants, which can be misleading *(13)*. Some studies have attempted to improve translation of PheWAS results using gene sets and pathways *(14)* or networks of PheWAS variants and diseases *(15, 16)*. However, these studies rely on the same variant-trait associations on which PheWAS are built and fall short of prioritizing likely actionable targets.

Integration of genomic, transcriptomic and other regulatory and functional information offers crucial justification for therapeutic target identification efforts, such as drug development *(17)*. Translational researchers also need access to this integrated information in a comprehensive platform that allows convenient investigation of complex relationships across multiple genes and traits. To meet this need, we present PhenomeXcan, a massive integrated resource of gene-trait associations to facilitate and support translational hypotheses. Predicted transcriptome association methods test the mediating role of gene expression variation in complex traits and organize variant-trait associations into gene-trait associations supported by functional information *(18–20).* These methods can describe direction of gene effects on traits, supporting how up- or down-regulation may link to clinical presentations or therapeutic effects. We trained transcriptome-wide gene expression models for 49 tissues using the latest Genotype-Tissue Expression data (GTEx; v8) *(21)* and tested the predicted effects of 8.87 million variants across 22,255 genes and 4,091 traits using an adaptation of the PrediXcan method *(18)*, Summary-MultiXcan, that uses summary statistics and aggregates results across tissues *(22)*. We then prioritized genes with likely causal contributions to traits using colocalization analysis *(23).* To make computation feasible given the large scale of data in this study, we developed fastENLOC, a novel Bayesian hierarchical colocalization method (see Methods). PhenomeXcan is the first massive gene-based (rather than variant-based) trait association resource. Our approach not only employs state-of-the-art techniques available to biologically prioritize genes with possible contributions to traits, but also presents information regarding pleiotropy and polygenicity across all human genes in an accessible way for researchers. Below, we provide several examples that showcase the translational relevance and discovery potential that PhenomeXcan offers.

## Results

### PhenomeXcan design and overall findings

We built a massive gene-to-phenome association resource that integrates GWAS results with gene expression and regulation data. We ran a version of PrediXcan *(18)*, Summary-MultiXcan, designed to use summary statistics and aggregate effects across tissues *(22)* on publicly available GWAS. In total, we tested the predicted effects of 8.87 million variants across 22,255 genes and 4,091 traits. Traits incorporate binary, categorical or continuous data types and range from basic anthropometric measurements to clinical traits and biochemical markers. We inferred association statistics (p-values and Z-scores) between predicted gene-expression variation and traits using optimal prediction models trained using 49 tissues from GTEx v8 *(21, 24, 25).* Non-causal, spurious gene-trait associations may be caused by linkage disequilibrium (LD) contamination and weighting of expression quantitative trait loci (eQTLs) *(21, 26)*. We therefore first performed Bayesian fine-mapping using the DAP-1/fgwas algorithm in TORUS *(27, 28)*. We then calculated the posterior probability of colocalization between GWAS loci and cis-eQTLs to prioritize possible causal genes via fastENLOC, a newly developed Bayesian hierarchical method that uses pre-computed signal clusters constructed from fine-mapping of eQTL and GWAS data to speed up colocalization calculations. The result is a matrix of 4,091 traits and 22,255 genes in which each intersection contains a PrediXcan p-value aggregated across 49 tissues and refined by a locus regional colocalization probability (locus RCP) (Figure 1). While a given colocalization threshold may be arbitrary, to minimize false negatives given the conservative nature of colocalization approaches *(26)*, we defined putative causal gene contributors as those genes with locus RCP > 0.1.

**Fig. 1:**
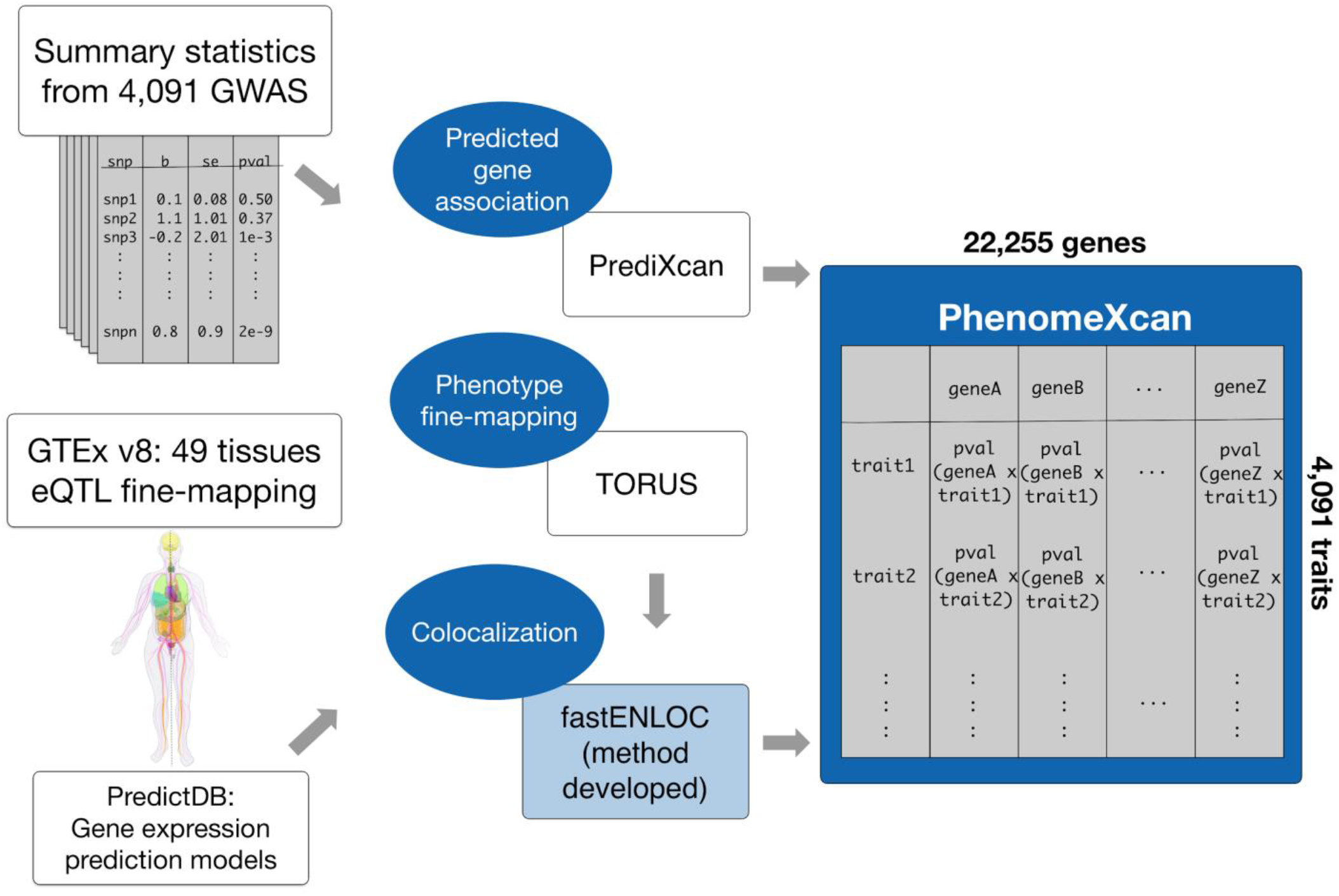
Schematic for the development of PhenomeXcan, a massive gene-based resource of gene-trait associations that can be used for translational hypothesis generation. Blue areas highlight methods we performed for this project, with fastENLOC being a novel colocalization method developed in the context of PhenomeXcan development. We developed PhenomeXcan by integrating genome-wide association study (GWAS) summary statistics with Genotype-Tissue Expression data (GTEx; v8) using PrediXcan methodology, then performing fine mapping and colocalization to identify the most likely causal genes for a given trait. PhenomeXcan is a massive resource containing PrediXcan p-values across 4,091 traits and 22,255 genes, aggregated across 49 tissues and refined by locus regional colocalization probability. (We thank Mariya Khan for the human illustration from the GTEx consortium.)

We found 65,603 significant associations (Bonferroni-corrected p-value < 5.5 × 10^−10^) across the entire genome/phenome space, where 19,579 (29.8 percent) had locus RCP > 0.1 (Supplementary Table S1). We constructed a quantile-quantile plot of all associations, which did not show evidence of systematic inflation (Supplementary Figure S1). These associations represent numerous potential targets for translational studies with biological support.

### Replicating known gene-trait associations

We evaluated PhenomeXcan’s performance using two different, independent validation approaches. For the first validation, we compared significant results from PhenomeXcan to significant results from the PheWAS Catalog, which combines the NHGRI-EBI GWAS catalog (as of 4/17/2012) and Vanderbilt University’s electronic health record to establish unique associations between 3,144 variants and 1,358 traits (https://phewascatalog.org/phewas) *(12, 29)*. We mapped traits from PhenomeXcan to those in the PheWAS Catalog using the Human Phenotype Ontology *(30)*. After filtering for genes included in both PhenomeXcan and the PheWAS Catalog, we tested 2,204 gene-trait associations. At a nominal p-value (p-value < 0.01), 1,005 PhenomeXcan gene-trait associations replicated with matched traits in the PheWAS catalog (AUC = 0.61; Figure 2A). Considering different methods of gene assignments for each GWAS locus (PheWAS: proximity, PhenomeXcan: Bayesian colocalization), we further evaluated our replication rate using random classifiers in a precision-recall curve (Figure 2B) and found considerable replicability between PhenomeXcan and PheWAS approaches compared to the null of no replication (α = 0.01, p-value < 1 × 10^−30^).

**Fig. 2:**
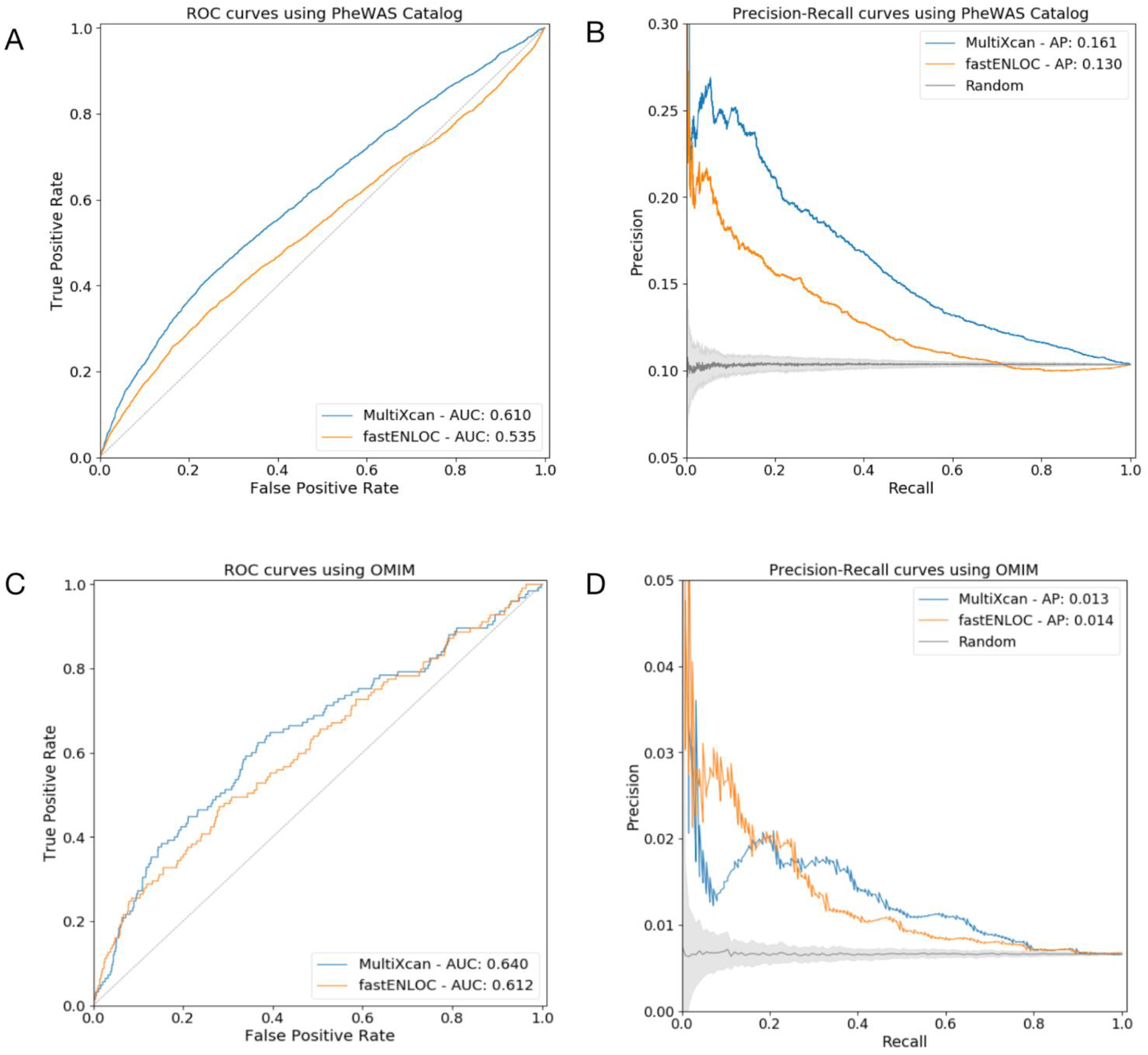
PhenomeXcan validation across the PheWAS Catalog and OMIM data sets using receiver-operating curves (ROC) and precision-recall (PR) curves. MultiXcan refers to the version of PrediXcan designed to take GWAS summary statistics and aggregate results across tissues *(22)*. **(A, B)** ROC curve and PR curve of PrediXcan significance scores (blue) and fastENLOC (orange) to predict PheWAS catalog gene-trait associations. **(C, D)** ROC curve and PR curve of PrediXcan significance scores (blue) and fastENLOC (orange) to predict OMIM catalog gene-trait associations. AUC refers to the area under the curve, AP refers to average precision. The predictive ability of both PrediXcan and fastENLOC demonstrate the statistical validity of PhenomeXcan associations.

For the second validation, we identified a set of high-confidence gene-trait associations using the Online Mendelian Inheritance in Man (OMIM) catalog *(31)*. We previously demonstrated that integrated analysis using PrediXcan *(18)* and colocalization *(23)* successfully predicts OMIM genes for matched traits *(26)*. We mapped 107 traits from PhenomeXcan to those in OMIM using the Human Phenotype Ontology *(30)* and curated a list of 7,809 gene-trait associations with support for causality. We compared gene-trait associations from this standard near GWAS loci (Supplementary Table S2) and found that PhenomeXcan successfully predicts OMIM genes (AUC = 0.64; Figure 2C). The limited precision seen here is expected in the setting of genes, such as those in OMIM, with large effects and rare variants (Figure 2D).

Of note, we did not filter any results by fastENLOC for either validation approach. The conservative nature of colocalization analysis can lead to increased false negatives *(26)*, which may contribute to decreased performance of fastENLOC in these scenarios.

### Identifying novel and underreported gene-trait associations

PhenomeXcan provides a resource for hypothesis generation using gene-trait associations, with over 19,000 potentially causal associations (p-value < 5.5 × 10^−10^, locus RCP > 0.1; Supplementary Table S1). As case studies, we discuss associations identified based on trait (“Morning/evening person (chronotype)”) and gene (*TPO*).

We reviewed the 15 most significant genes associated with “Morning/evening person (chronotype)” (a UK Biobank trait) based on PrediXcan p-values across the 49 tissues and locus RCP > 0.1 (Supplementary Table S3). Three of 15 genes had not been previously reported in any GWAS involving UK Biobank subjects related to sleep or chronotype: *VIP, RP11-220I1.5* and *RASL10B*. Notably, a variant associated with *VIP* (p-value=1.812 × 10^−17^, locus RCP=0.30) is discussed in a GWAS of 89,283 individuals from the 23andMe cohort who self-report as “a morning person” (rs9479402 near *VIP*, 23andMe GWAS p-value=3.9 × 10^−11^) *(32)*. *VIP* produces vasoactive intestinal peptide, a neurotransmitter in the suprachiasmatic nucleus associated with synchronization of circadian rhythms to light cycles *(33)*. The long noncoding RNA *RP11-220I1.5* (p-value=6.427 × 10^−11^, locus RCP=0.22) and the gene *RASL10B* (p-value=1.098 × 10^−10^, locus RCP=0.17) have not been previously reported in any GWAS or functional/clinical studies associated with this trait. *RASL10B* produces a 23 kiloDalton GTPase protein that demonstrates overexpression in the basal ganglia in GTEx *(21)*, potentially representing a novel association. Besides *VIP*, three other genes in this set had clinical/functional studies associated with sleep or chronotype in PubMed: *RAS4B, CLN5* and *FBXL3*. *RAS4B* (p-value=1.660 × 10^−19^, locus RCP=0.64) was linked to a transcriptional network regulated by *LHX1* involved in circadian control *(34)*. *CLN5 (*p-value=5.248 × 10^−18^, locus RCP=0.37) mutations are associated with neuronal ceroid lipofuscinosis, which can manifest with sleep-specific dysfunction *(35)*. *FBXL3 (*p-value=1.54 × 10^−16^, locus RCP=0.41) assists with turnover of the *CRY* protein through direct interaction to regulate circadian rhythms *(36).* Our results also note *VAMP3* (p-value=7.317 × 10^−18^, locus RCP=0.67), a gene with little research in chronotype or sleep, which lies adjacent to *PER3. PER3* is one of the *Period* genes characterized as part of the circadian clock and described in numerous functional studies, animal models and human polymorphism association studies *(37)*. Both *VAMP3* and *PER3* (p-value=1.65 × 10^−17^) are significant in PhenomeXcan, with *PER3* showing a lower level of colocalization with locus RCP=0.1. PhenomeXcan, to our knowledge, is one of the first hypothesis-generating tools to provide unbiased links between a trait and associated genes for the researcher’s evaluation. In conjunction to rich knowledge obtained from functional studies, PhenomeXcan can be used to generate or support subsequent translational efforts.

We next evaluate PhenomeXcan as a platform to study novel and underreported gene-trait associations. Thyroid peroxidase (*TPO*) encodes a membrane-bound glycoprotein that plays a crucial role in thyroid gland function *(38)*. The strongest associations in PhenomeXcan support the known role of *TPO* in thyroid hormone production: “Self-reported hypothyroidism or myxedema” (p-value=1.40× 10^−14^, locus RCP=0.99) and “Treatment with levothyroxine” (p-value=1.54× 10^−10^, locus RCP=0.99). Hypothyroidism has been clinically linked to increased respiratory symptoms. Although the mechanism for this is not well understood *(39)*, our results suggest that these could be explained by common genetic factors; “Treatment with salmeterol” (a medication used to treat lung disease such as asthma or chronic obstructive pulmonary disease) showed moderate associations with TPO in PhenomeXcan (p-value=7.45× 10^−5^, locus RCP < 0.1). *TPO* is also contained in the NIH Biosystems Pathways for the development of pulmonary dendritic cells *(40)*. “Time to complete round” (drawing as a measure of cognitive function) showed another moderate association in PhenomeXcan (p-value=1.19× 10^−4^, locus RCP < 0.1). Thyroid function has been clinically linked to time to draw a clock as a form of cognitive measurement *(41)*. Other trait associations identified in PhenomeXcan with *TPO* include “Single major depression episode” (p-value=2.48× 10^−4^, locus RCP < 0.1) and “Treatment with doxazosin” (a medication used in the UK for hypertension) (p-value=8.80 × 10^−4^, locus RCP=0.12), both of which have demonstrated clinical association with thyroid abnormalities *(42,43)*. To our knowledge, none of these traits have been deeply investigated with *TPO* previously, highlighting how PhenomeXcan may be useful in expanding gene-trait association studies and functional studies through consideration of independent traits associated with a given gene.

### Revealing complex clusters of pleiotropy and polygenicity for translational hypotheses

PhenomeXcan allows more complex exploration of associated genes and traits beyond individual queries. As an example, to study genes associated with white blood cell count, we can cluster related genes and traits. Starting from the trait “Lymphocyte percentage,” the top associated genes include *PSMD3*, *CD69, KLF2, CXCL2, CREB5, CXCL3, ZFP36L2, JAZF1, NCOR1*, and *TET2*. These genes represent pathways associated with chemokine and interleukin signaling as well as peptide ligand binding, but are not specific to one particular pathway or genomic location *(44).* We can assess these genes’ associations with white blood cell traits (neutrophil count/percentage, lymphocyte count/percentage, eosinophil count/percentage, monocyte and basophil percentages) and infer some understanding of their causal mechanism. *PSMD3*, for instance, demonstrates stronger associations with neutrophil and lymphocyte traits (mean p-value < 1× 10^−30^, mean locus RCP=0.43), whereas *ZFP36L2* demonstrates consistent associations across white blood cell, platelets and red blood cell traits (mean p-value < 1.54 × 10^−24^, mean locus RCP=0.27) (Figure 3). Disruption of *ZFP36L2* results in defective hematopoiesis in mice *(45)*, whereas *PSMD3* has been identified in genome-wide association studies related to white blood cell count and inflammatory states *(46).* Clusters of associated genes and traits can support more robust translational hypotheses through similarities in associations and generate more nuanced experimental designs through differences between associations.

**Fig. 3:**
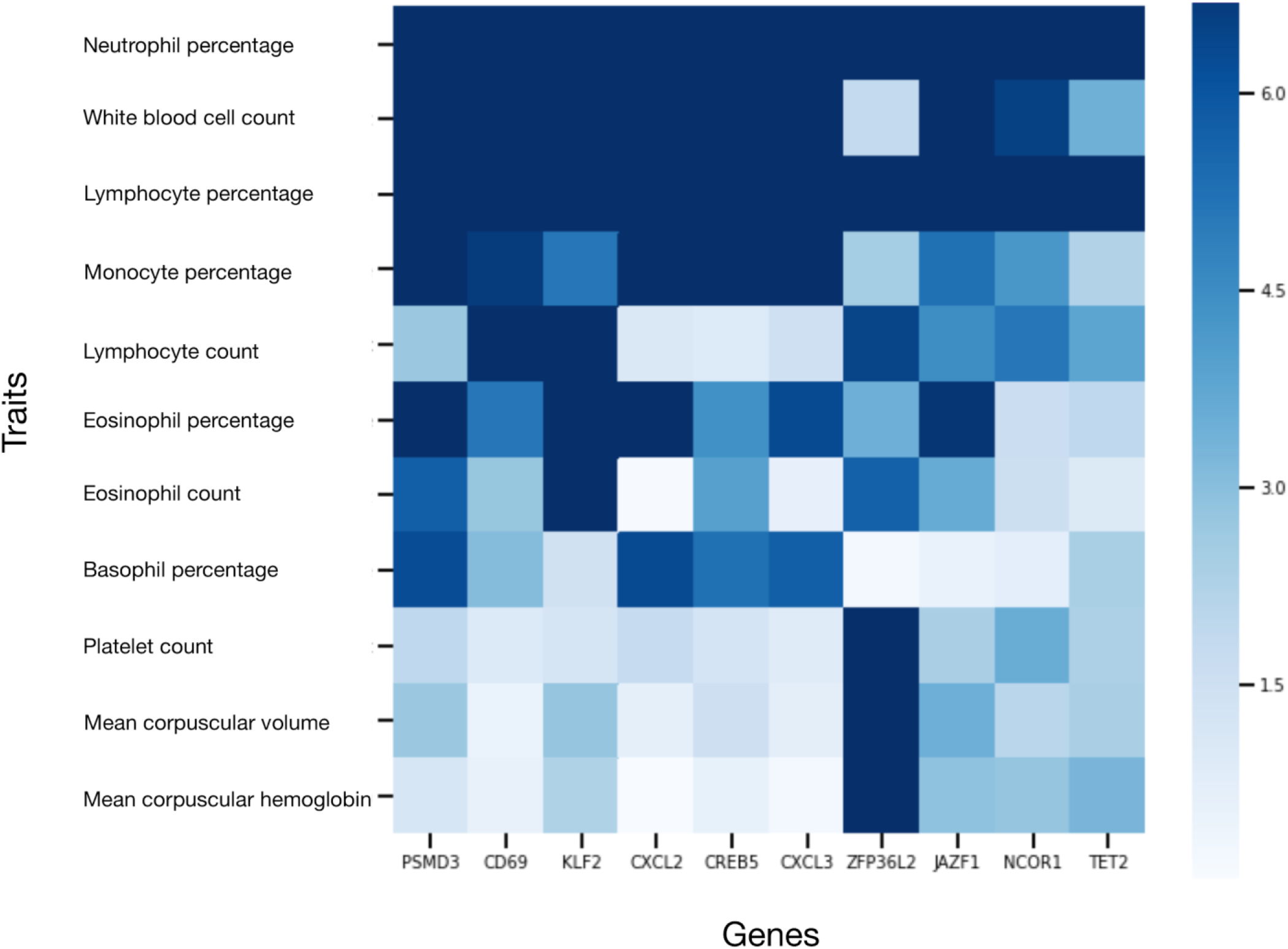
Visual heatmap cluster of gene-trait associations for white blood cell traits identified in PhenomeXcan. Z-scores are derived from PrediXcan p-values, with the ceiling of association (dark blue) > or equal to 7. In this heatmap, we demonstrate the associations between the genes *PSMD3*, *CD69, KLF2, CXCL2, CREB5, CXCL3, ZFP36L2, JAZF1, NCOR1*, and *TET2* and the white blood cell traits “Neutrophil count” and “Neutrophil percentage”, “Lymphocyte count” and “Lymphocyte percentage”, “Eosinophil count” and “Eosinophil percentage”, “Monocyte percentage” and “Basophil percentage.” “Platelet count” and “mean corpuscular volume” (for red blood cells) serve as alternate blood traits. T*ZFP36L2* has consistent associations across platelets and red blood cells relative to other genes. Accordingly, functional studies demonstrate *ZFP36L2* plays a role in hematopoiesis, whereas studies support the others genes’ involvement in inflammation-related pathways or diseases. These types of clusters can support hypotheses and experimental designs regarding the mechanisms through which genes contribute to traits.

### Discovering links between common traits and rare diseases

PhenomeXcan can also be integrated with any gene-trait databases to explore pleiotropically linked traits and shared associated genes. We integrated PhenomeXcan with ClinVar, a publicly available archive of rare human diseases and associated genes (including OMIM) and one of the most widely used gene-trait databases in the clinical setting *(47)*. We examined the associations between the 4,091 GWAS-derived traits in PhenomeXcan and 5,094 ClinVar diseases by (a) calculating PrediXcan Z-scores for every gene-trait association in PhenomeXcan and (b) for each PhenomeXcan/ClinVar trait pair, we computed the average squared PrediXcan Z-score considering the genes reported in the ClinVar trait (see Methods). We then created a matrix of PhenomeXcan traits by ClinVar traits with mean squared Z-scores (Figure 4A, Figure 4B), where peaks represent shared genes. We defined significant associations between traits as those with Z-score > 6; this represents the equivalent of a Bonferroni-adjusted p-value of 0.05 based on our map of the distribution of Z-scores (Supplementary Figure S2).

**Fig. 4:**
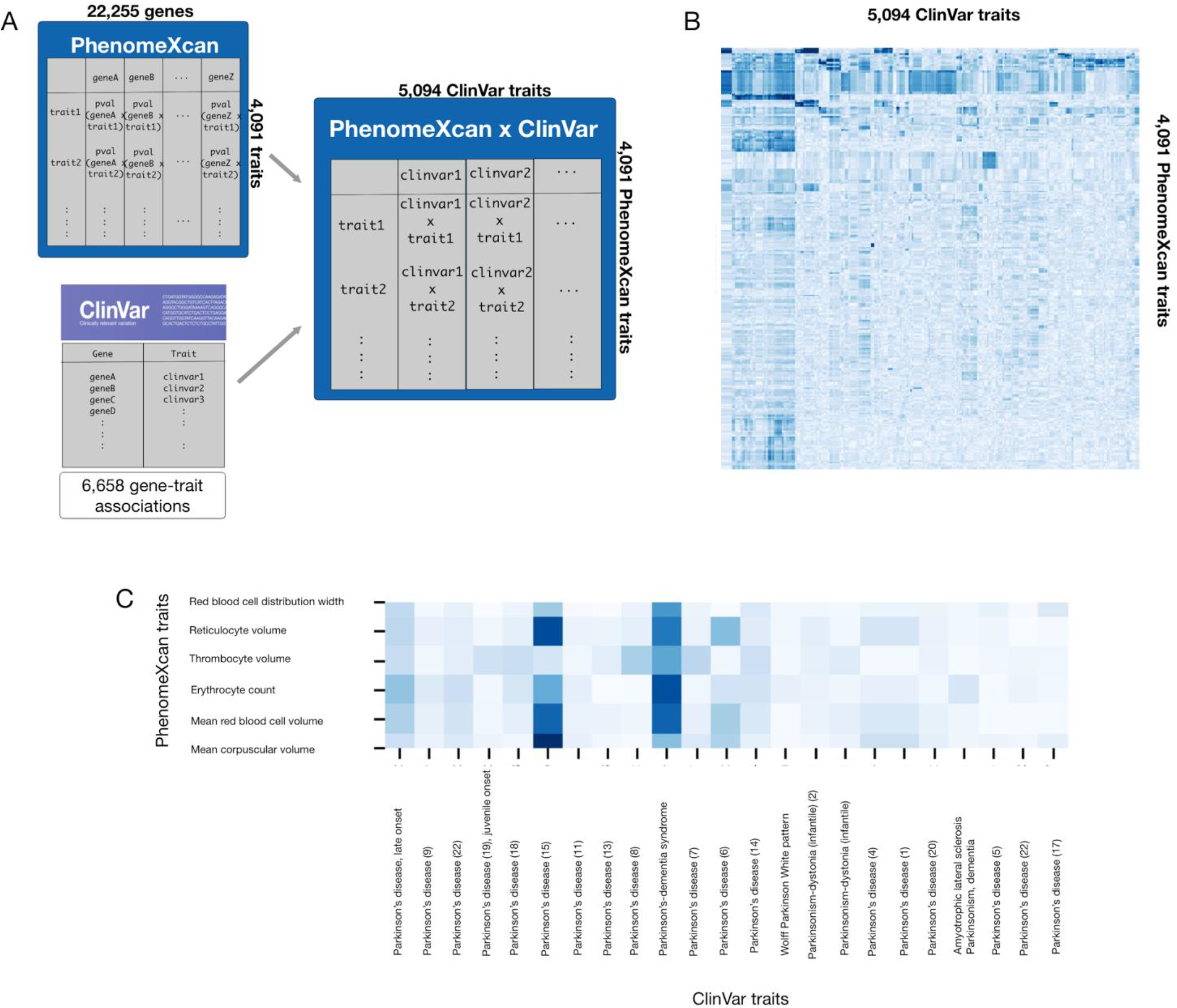
Schematic and visualization of PhenomeXcan x ClinVar. **(A)** Schematic depicting the development of PhenomeXcan x ClinVar. For each PhenomeXcan/ClinVar trait pair, we computed the average squared PrediXcan Z-score considering the genes reported in the ClinVar trait. **(B)** Heatmap visualizing associations in PhenomeXcan x ClinVar. Darker blue represents stronger association. Again, complex clusters of inter-trait associations can be identified to link common traits and rare diseases. **(C)** Heatmap demonstrating linked traits in PhenomeXcan (rows) and ClinVar (columns) for the example association between Parkinson’s disease and red blood cell traits. We see the strongest associations between mean corpuscular volume, mean reticulocyte volume and mean spherical red cell volume and “Parkinson disease 15.” In ClinVar, each variant of Parkinson’s disease linked to a different gene is listed under a different number, making it unsurprising that associations to other forms of Parkinson’s disease are not as strong.

As an example, we found links between the ClinVar trait “Parkinson disease 15” and the following traits: mean corpuscular volume, mean reticulocyte volume and mean spherical red cell volume (Figure 4C). The driving gene for these blood traits linked to “Parkinson disease 15” was *FBX07* (mean Z-score across all traits=20.4. mean locus RCP=0.968). *FBX07* plays a role in the ubiquitin system linked to Parkinson’s disease *(48)*. Two GWAS (the HaemGen consortium and eMERGE) link *FBX*07 with mean corpuscular volume *(49,50).* Through PhenomeXcan, we discover a pleiotropic relationship between Parkinson’s disease and red blood cell traits mediated through *FBX07* that has not been studied in humans. Validating this finding, a mouse model has been designed specifically to study this pleiotropic effect *(51)*. This case study demonstrates how this powerful variation on PhenomeXcan can significantly improve translational hypothesis generation by supporting genetic links between associated rare diseases and common traits across research platforms.

### Identification of potential therapeutic drug targets and related adverse effects

PhenomeXcan offers direct translational applicability, providing genomic evidence to support therapeutic targets and associated side effects. As an example, *PCSK9* is a genetically supported, clinically validated target for cardiac prevention through inhibition of its binding to the LDL receptor and reduction of blood LDL cholesterol levels *(52)*. We can study the cluster of genes and traits produced by *PCSK9* in PhenomeXcan for relevant information about this target. Most of the traits with strongest associations to *PCSK9* relate to diagnosis and treatment of elevated cholesterol or atherosclerosis, including familial heart disease. Because inherited *PCSK9* variation is associated with increased likelihood of type 2 diabetes, there was concern that *PCSK9* therapies could elevate risk to type 2 diabetes. The inhibiting drugs therefore required large substudies from clinical trials to confirm no association with worse diabetes *(53,54)*. While not at genome-wide significance, *PCSK9* associates with type I diabetes in PhenomeXcan (p-value=3.88 × 10^−4^, locus RCP<0.1). We recognize that type I and type 2 diabetes have different clinical etiologies. For the purpose of drug development, though, assessing *PCSK9* in PhenomeXcan produces both its primary target (blood cholesterol levels as related to atherosclerosis) and, through independently identified traits, potential adverse effects via diabetes. The most commonly represented genes associated with the strongest traits for *PCSK9* include *APOE, LDLR, APOB, PSRC1, CELSR2, SORT1, ABCG8, ABCG5, and HMGCOR*. Unsurprisingly, all of these genes have all been implicated in genetic susceptibility to hypercholesterolemia (some, such as *SORT1*, may be the primary causative gene in their pathway) *(55)*. Examining potential targets in PhenomeXcan could not only help anticipate side effects via independent traits, but also identify related gene networks / alternative targets with therapeutic relevance.

## Discussion

In this paper, we introduce PhenomeXcan, an innovative, powerful resource that makes comprehensive gene-trait associations easily accessible for hypothesis generation. Using PrediXcan allows us to derive gene-based associations with traits in context by integrating GWAS summary statistics with transcriptome-wide predicted expression and regulatory / functional information. We previously demonstrated that integrated analysis using PrediXcan and colocalization improves precision and power for target gene identification *(26)*. To build PhenomeXcan, we also develop a novel, rapid colocalization method, fastENLOC, that could handle data at this scale (4,091 traits × 22,255 genes × 49 tissues) (see Methods). PhenomeXcan implements the best practices derived from applying GTEx v8 *(21, 25)* to biologically prioritize genes with possible causal contribution to a given trait.

PhenomeXcan’s flexible structure and adaptability allow translational researchers to easily explore clinically relevant questions. The resource can be queried by gene or trait and allows identification of novel and underrepresented associations. It offers exploration of polygenicity and pleiotropy dimensions by allowing for queries across multiple genes and traits. It can also be integrated with other gene-trait datasets to explore linked traits and report common associated genes. We offer ClinVar as an example, but any deeply annotated database of genes and traits may be integrated in this manner. Other possible translational uses of PhenomeXcan include biomarker exploration, identification of clinically relevant disease modifiers, and polygenic score building (using genes associated with queried traits), as well as novel directions for basic science collaborations and clinical study of linked traits (using traits associated with queried genes).

We note some caveats. Diseases with variability not related to changes in gene expression (e.g. epigenetic regulation or traits with important environmental contributions) are not expected to be captured well by this method. Our model also better captures common overall genetic contributors rather than genes identified from rare variants. We do note that our ClinVar validation standard tends to favor larger-effect genes with monogenic etiology, while the PhenomeXcan association method itself is less biased. Regulatory pleiotropy is widespread across the genome *(21)*. In our chronotype example, *VAMP3* and *PER3* demonstrate regulatory pleiotropy. With that degree of proximity, large-scale tools are not able to distinguish causal genes well *(21)*. We provide this example to acknowledge how PhenomeXcan encounters this phenomenon and show the benefit of performing these associations across all human genes. We offer colocalization as a possible means of prioritizing causal variants, but both significance of association and colocalization must be taken into account in our results. Work from large-scale statistical genetics tools, such as PhenomeXcan, and Mendelian genetics / functional studies must then be combined in order to best understand the breadth of genetic contributors to complex traits. We have favored a locus RCP threshold of 0.1 to limit false negatives related to colocalization. Poor regional colocalization probability (locus RCP~0) may reflect a lack of sufficient evidence with available data, particularly for understudied genes, rather than true lack of causality. We therefore reported traits in this paper that had a locus RCP < 0.1, but had functional support for potential association. Similarly, the genome-wide threshold of significance is conservative, and we discuss associations with functional support even with less significant p-values. Importantly, GWAS summary statistics used in this project were for subjects and patients of European ancestry. Improving the applicability of this type of work to global populations remains of paramount importance throughout genetic medicine, and we will continue to integrate more GWAS summary statistics from broader consortia.

Resources that translate biologically relevant genomic and transcriptomic information into gene-trait associations are already critical for hypothesis generation and clinically relevant research *(56).* We offer PhenomeXcan, an integrated mapping for the function of every human gene, as a publicly available resource to advance the investigation of complex human diseases by improving the accessibility of relevant links between the entire genome and the phenome.

## Materials and Methods

### Trait selection and preprocessing/quality control of variants

We developed PhenomeXcan with 4,091 traits from publicly available GWAS summary statistics. Summary statistics from GWAS performed for 4,049 traits from the UK Biobank (on 361,194 samples) were obtained from the publicly available dataset compiled by the Neale Lab at the Broad Institute (*57*); we did not use individual-level data. The UK Biobank is a prospective cohort of approximately 500,000 subjects between 40 and 69 years of age, recruited from 2006-2010 in the United Kingdom *(58)*. Traits characterized by the Neale lab include 2,891 auto-curated traits using PHESANT (*59*), of which 274 are continuous, 271 ordinal and 2,346 binary. 633 binary traits were extracted from hospital-level data (ICD-10 codes). 559 traits were manually curated in collaboration with the FinnGen Consortium. Traits available cover a range of categories, from lifestyle traits and socio-demographic questions to clinical biomarkers and diagnoses. Separate sex-specific summary statistics and sex chromosome analyses were not included in this project. More details on the GWAS derivations and quality control is provided in the website of the project: http://www.nealelab.is/uk-biobank. We do note that for these GWAS, 361,194 individuals were selected for inclusion based on quality of genotypes, white British ancestry (based on both self-report and principal components analysis). Only those variants with an imputation quality score (INFO) > 80%, a minor allele frequency (MAF) > 0.1%, call rate > 95% and a Hardy-Weinberg equilibrium p-value > 1 × 10^−10^ were selected.

We also compiled 42 additional traits from summary statistics from publicly available GWAS and GWAS-meta analyses external to the UK Biobank study both to validate synthesis of additional GWAS data and to overcome limitations related to poor sample sizes in the UK Biobank for specific diseases (e.g. breast cancer). These GWAS and traits represent a broad array of disease-related categories, including immunological response, psychiatric and neurologic traits, cardiometabolic diseases and syndromes and cancer. We have previously described the harmonization and imputation process *(26)* (Supplementary Table S4).

ClinVar is a publicly available archive of clinically reported human genetic variants and associations with disease maintained by the National Institutes of Health (https://www.ncbi.nlm.nih.gov/clinvar/). Variant associations with disease are identified by manual review of submitted interpretations from “clinical testing laboratories, research laboratories, locus-specific databases, Online Mendelian Inheritance of Man (OMIM), GeneReviews, UniProt, expert panels and practice guidelines” *(31, 47).* Traits can be reported to ClinVar as a single concept or set of clinical features. When possible, traits are mapped manually to standardized terms from databases including OMIM and the Human Phenotype Ontology (HPO) *(30)*. All gene-trait associations published by ClinVar for 7/2019 were used for integration with PhenomeXcan.

### PrediXcan and Summary-MultiXcan (S-MultiXcan)

S-MultiXcan is a method in the PrediXcan family *(18)* that associates genes and traits by testing the mediating role of gene expression variation in complex traits, but (a) requires only GWAS summary statistics and (b) uses multivariate regression to combine expression information across tissues *(22).* First, linear prediction models of genotype in the vicinity of the gene to expression are trained in reference transcriptome datasets such as the Genotype-Tissue Expression project (GTEx) *(21)*. Second, predicted expression based on actual genetic variation is correlated to the trait of interest to produce a gene-level association result for each tissue. In S-MultiXcan, the predicted expression is a multivariate regression of expression across multiple tissues. In order to avoid collinearity issues and numerical instability, the model decomposes the predicted expression matrix into principal components and keeps only the eigenvectors of non-negligible variance. We considered a PCA regularization threshold of 30 to be a conservative choice. This approach improves detection of associations relative to use of one tissue type alone and offers a reduced false negative rate relative to a Bonferroni correction. We used optimal prediction models based on the number and proportion of colocalized gene level associations *(26)*. These models select features based on fine-mapping *(24)* and weights using expression quantitative trait loci (eQTL) effect sizes smoothed across tissues using mashr *(25)*. The result of this approach is a genome-wide gene-trait association list for a given trait and GWAS summary statistic set.

### Colocalization of GWAS and eQTL signals

Bayesian fine-mapping was performed using TORUS (*28*). We estimated probabilities of colocalization between GWAS and cis-eQTL signals using Bayesian regional colocalization probability, as performed in the ENLOC methodology (*23*). For this particular study, given the large scale of the data, we developed a novel implementation, entitled fastENLOC.

### fastENLOC

fastENLOC is a novel method we developed that combines the speed of eCAVIAR *(60)* and the biological factors incorporated into ENLOC *(23)*. eCAVIAR assumes that the probability of a variant being causal for a trait is independent of the probability of the variant causally affecting gene expression, which results in rapid processing but can be too conservative. ENLOC, by contrast, requires significant processing time but estimates biological dependence and colocalization priors using eQTL enrichments among GWAS signals.

fastENLOC takes advantage of Bayesian signal clusters (or credible sets) constructed from fine-mapping analysis of eQTL and GWAS data and provides improved precision for colocalization analysis. Signal clusters consist of variants in linkage disequilibrium (LD) and serve as natural analytic units for colocalization analysis, representing the same underlying independent association signals. fastENLOC automatically assesses a locus-specific regional colocalization probability (locus RCP) for each signal cluster inferred from eQTL analysis. As with eCAVIAR, fastENLOC also allows direct input of posterior inclusion probabilities from GWAS analysis, enabling colocalization of multiple potential association signals from a single GWAS locus.

fastENLOC is implemented in a self-contained C++ program and runs magnitude faster than ENLOC. Despite their different approaches to colocalization analyses, fastENLOC and ENLOC agree with locus RCP reporting (Supplementary Figure S3).

The software and its source code are freely available on Github at http://github.com/xqwen/fastenloc/.

We provide a brief derivation of its approach: Let ***D*, *E*** denote the association data from GWAS and eQTL analyses, respectively. Let 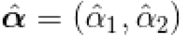 denote the point estimate of the enrichment vector. We consider a signal cluster inferred from the fine-mapping analysis of either eQTLs or GWAS and use latent binary indicator *p*-vectors ***d*, γ** to represent the causal association status of its *p* member single-nucleotide polymorphisms (SNPs) with the complex trait and the gene expression level of interest, respectively.

A signal cluster, by definition, contains a set of SNPs in LD and represent the same underlying genetic association signal. Furthermore, we use γ_0_ to denote the configuration of no causal eQTLs in the cluster and γ_1_ to denote the *i*th SNP is the true causal eQTL SNP (i.e., the *i*th entry is set to 1 and 0 for the remaining SNPs).

Assuming GWAS data are originally analyzed using an exchangeable prior 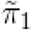, i.e.,

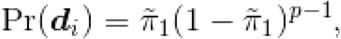

and

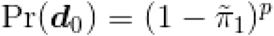

By the nature of a signal cluster, it follows from the Bayes rule that

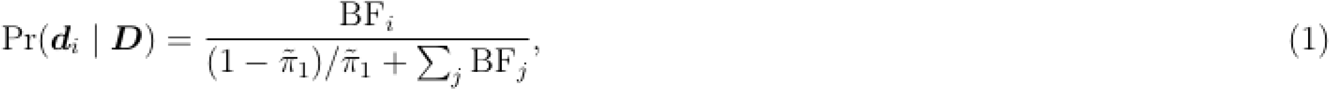

where **BF**_*i*_ denotes the marginal likelihood ratio,

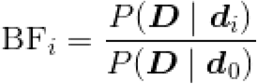

Note that in case that the GWAS posterior probability is derived from a multi-SNP analysis, **BF**_*i*_ may not be well-approximated by single SNP testing statistics. Nevertheless, given 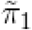 and note that Pr(γ_*i*_ | ***D***) coincides with the posterior inclusion probability (PIP) of the *i*th SNP in the signal cluster, **BF**_*i*_’s can be straightforwardly computed from equation (1). Additionally, 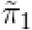 can be obtained by averaging the PIPs from all interrogated SNPs.

Given the enrichment information, the GWAS prior differs for eQTL and non-eQTL SNPs. Specifically, for eQTL SNP,

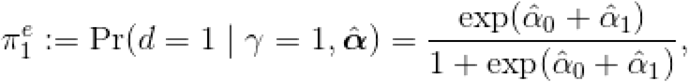

and for non-eQTL SNP,

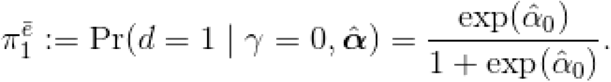

Using the eQTL-informed priors, the GWAS posterior probability can be updated analytically, i.e.,

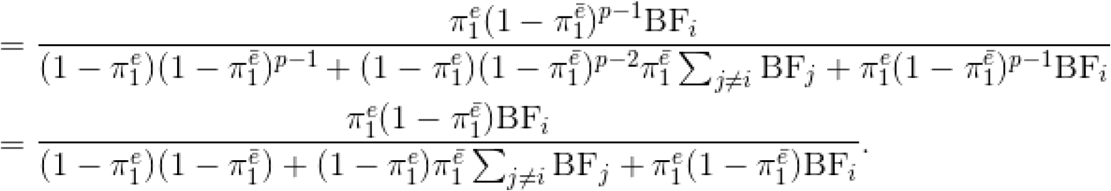

Subsequently, the colocalization probability at the *i*th SNP is computed by

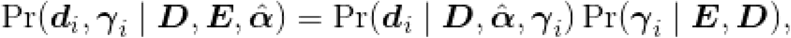

where we approximate Pr(γ_*i*_ | ***E, D***) with the eQTL PIP for the *i*th SNP. The regional colocalization probability, RCP, for the signal cluster of interest is given by

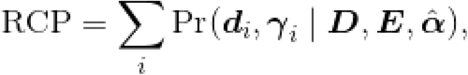

because events {γ_*i*_, ***d***_*i*_} and {γ_*j*_, ***d***_*j*_} for *i* ≠ *j* are mutually exclusive within a signal cluster.

### Validation of PhenomeXcan using PheWAS and ClinVar

We evaluated the accuracy of gene-trait associations in PhenomeXcan by using two different gene-trait association datasets and deriving the receiver-operator (ROC) and precision-recall (PR) curves for each. We mapped traits from PhenomeXcan to those in either PheWAS Catalog *(29)* or OMIM *(31)* by using the Human Phenotype Ontology (*30*) and the GWAS Catalog as intermediates. For traits in the PheWAS Catalog, we tested 2,204 gene-trait associations that could be mapped in both PhenomeXcan and the PheWAS Catalog, from a total 21,323 gene-traits associations consisting of all genes present in an LD block with GWAS signal. For the OMIM traits, we developed a standard (Supplementary Table S2) of 7,809 high-confidence gene-trait associations that could be used to measure the performance of PhenomeXcan, of which 125 could be mapped to GWAS loci. This standard was obtained from a curated set of trait-gene pairs from the OMIM database by mapping traits in PhenomeXcan to those in OMIM (*31*). Briefly, traits in PhenomeXcan were mapped to the closest phecode using the GWAS catalog-to-phecode. Then we created a map from phecodes to terms in the Human Phenotype Ontology (HPO), which allowed us to link our GWAS traits to OMIM disease description by utilizing phecodes and HPO terms as intermediate steps. For each gene-trait pair considered causal in this standard, we determined if PhenomeXcan identified that association as significant based on the resulting p-value. We did not filter results based on locus RCP in these validations to avoid worsened performance due to false negatives.

### Supporting evidence for PhenomeXcan results

PhenomeXcan results for case studies were included based on their p-values and locus RCP. We defined putative causal gene contributors as those genes with p-values less than 5.5 × 10^−10^ and locus RCP > 0.1. Given these conservative measures, however, we did discuss associations that were less significant or had a lower locus RCP with functional evidence. We used the NHGRI-EBI GWAS Catalog (10/21/2019) to identify GWAS results both using the UK Biobank (given the predominance of this dataset in PhenomeXcan) and other datasets. We performed systematic literature searches on PubMed using the gene name alone, with the specific trait category and trait name to identify functional studies relevant to a trait of interest.

### Building PhenomeXcan x ClinVar

We examined links between 4,091 PhenomeXcan traits and 5,094 ClinVar traits and associated genes. ClinVar traits were excluded if they did not have known associated genes in PhenomeXcan. To compare a PhenomeXcan trait *t* and a ClinVar trait *d*, we calculated the mean squared Z-score:

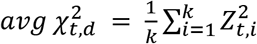

where *k* is the number of genes reported in ClinVar for trait *d*, and *Z* is the Z-score of gene *i* obtained with S-MultiXcan for trait *t.* We then created a matrix of PhenomeXcan traits by ClinVar traits with mean squared Z-scores. We defined significant associations between traits as those with Z-score > 6; this represents the equivalent of a Bonferroni-adjusted p-value of 0.05 based on our map of the distribution of Z-scores (Supplementary Figure S2).

## Acknowledgements

This research benefited from the use of credits from the National Institutes of Health (NIH) Cloud Credits Model Pilot, a component of the NIH Big Data to Knowledge (BD2K) program. We thank Julian Solway for helpful discussion and feedback on the manuscript.

## Funding

This work is supported by National Institutes of Health grants R01MH107666 (H.K.I.) and P30DK020595 (H.K.I.). This work was completed in part with computational resources provided by Bionimbus *(61)*, and the Center for Research Informatics. The Center for Research Informatics is funded by the Biological Sciences Division at the University of Chicago with additional funding provided by the Institute for Translational Medicine, CTSA grant number UL1 TR000430 from the National Institutes of Health. This work was also supported by funding from the Small Grants program from the University of Chicago Biological Sciences Division (BSD) Office of Diversity and Inclusion (M.D.P.) and the BSD Career Advancement for Postdocs Travel Award (M.D.P.).

## Author Contributions

All authors discussed the results and interpretation, and commented on the manuscript. In addition to the latter activities, Milton Pividori: Performed the large scale computation and other analyses, drafted the manuscript and prepared figures and tables. Padma Sheila Rajagopal: Performed analyses and drafted the manuscript. Yanyu Liang: Provided software and data for silver standard development and performance measurements. Alvaro Barbeira: Performed analysis, created database and web application for sharing of PhenomeXcan. Owen Melia: Supported database and web application development for PhenomeXcan. Lisa Bastarache: Assisted with construction of silver standard. YoSon Park: Edited the manuscript and provided insights. Xiaoquan Wen: Developed the theory and implemented fastENLOC in C++. Hae Kyung Im: Conceived PhenomeXcan, supervised the implementation and analysis, edited the manuscript. Authors report the following declarations: HKI reports speaker honoria received from GlaxoSmithKline and AbbVie.

## Data and materials availability

PhenomeXcan is publicly available at phenomexcan.org. The site contains the results of S-PrediXcan (individual tissues reported) and S-MultiXcan (across all tissues) applied to 4,091 traits and 22,255 genes. PhenomeXcan can be queried by gene (to result in traits) or trait (to result in genes). Multiple genes or traits can be queried at once. The result will list associations by p-value (from either S-PrediXcan if tissue-specific or S-MultiXcan as the best across tissues) and locus RCP from fastENLOC. We have also provided a queryable table of PhenomeXcan’s 4,091 traits × 5,094 ClinVar traits. Queries can be made by either PhenomeXcan trait or ClinVar trait, and the result will list associated traits, shared genes in the association and mean Z-score. The data sets used in this paper is publicly available in [Zenodo DOI]. Scripts to generate our results will be available on Github at https://github.com/hakyimlab/phenomexcan.

## Supplementary Materials

File name: suppl_table_S1-significant_gene_trait_associations.xlsx

**Supplementary Table S1: Gene-trait associations in PhenomeXcan that were significant by Bonferroni correction p-value and with locus RCP > 0.1**.

This table contains all **19,579 gene-trait associations** with p-value < 5.5 × 10^−10^ and locus RCP > 0.1.

File name: suppl_table_S2-UKBiobank_to_OMIM-standard.xlsx

**Supplementary Table S2: Standard of OMIM gene-trait associations used to validate PhenomeXcan**

This table contains 7,809 high-confidence gene-trait associations from OMIM that were used to evaluate the performance of PhenomeXcan.

**Supplementary Table S3:** Summary of genes and evidence associated with UK Biobank trait “Morning/evening person (chronotype)”. Genes are sorted by PrediXcan p-value for the best tissue expression, with locus regional colocalization probability (locus RCP) higher than 0.1. Higher p-values and locus RCP scores suggest greater likelihood of causal association to the trait. Evidence is organized by gene reports in GWAS using UK Biobank subjects, GWAS not using UK Biobank subjects, and clinical/functional studies. GWAS were identified using the NHGRI-EBI GWAS catalog (10/21/2019), and functional/clinical studies were identified from PubMed using searches for the gene name as well as the gene name/trait category and gene name/trait.

| **Gene** | **Chromosome** | **p-value** | **Locus RCP** | **Number of UK Biobank GWAS** | **Number of non-UK Biobank GWAS** | **Number of clinical or functional studies focused on sleep or chronotype mechanisms** |
| --- | --- | --- | --- | --- | --- | --- |
| *TRAF3IP1* | 2q37.3 | 1.724e-20 | 0.43 | 4 | 1 | 0 |
| *RASA4B* | 7q22.1 | 1.660e-19 | 0.64 | 1 | 0 | 1 |
| *CPNE8* | 12q12 | 3.231e-18 | 0.12 | 2 | 1 | 0 |
| *CLN5* | 13q22.3 | 5.248e-18 | 0.37 | 4 | 1 | 3 |
| *VAMP3* | 1p36.23 | 7.317e-18 | 0.67 | 0 | 1 | 0 |
| *VIP* | 6q25.2 | 1.812e-17 | 0.30 | 0 | 1 | 7 |
| *FBXL3* | 13q22.3 | 1.545e-16 | 0.41 | 4 | 1 | 29 |
| *TNRC6B* | 22q13.1 | 8.441e-14 | 0.19 | 6 | 1 | 0 |
| *RASD1* | 17p11.2 | 1.246e-12 | 0.23 | 4 | 1 | 0 |
| *ZCCHC7* | 9p13.2 | 4.282e-11 | 0.29 | 2 | 0 | 0 |
| *RP11-220I1.5* | 9 | 6.427e-11 | 0.22 | 0 | 0 | 0 |
| *EBLN3P* | 9p13.2 | 6.853e-11 | 0.95 | 2 | 0 | 0 |
| *RASL10B* | 17q12 | 1.098e-10 | 0.17 | 0 | 0 | 0 |
| *PMFBP1* | 16q22.2 | 1.413e-10 | 0.86 | 4 | 0 | 0 |
| *DDI2* | 1p36.21 | 2.156e-10 | 0.30 | 4 | 0 | 0 |

**Supplementary Table S4:**
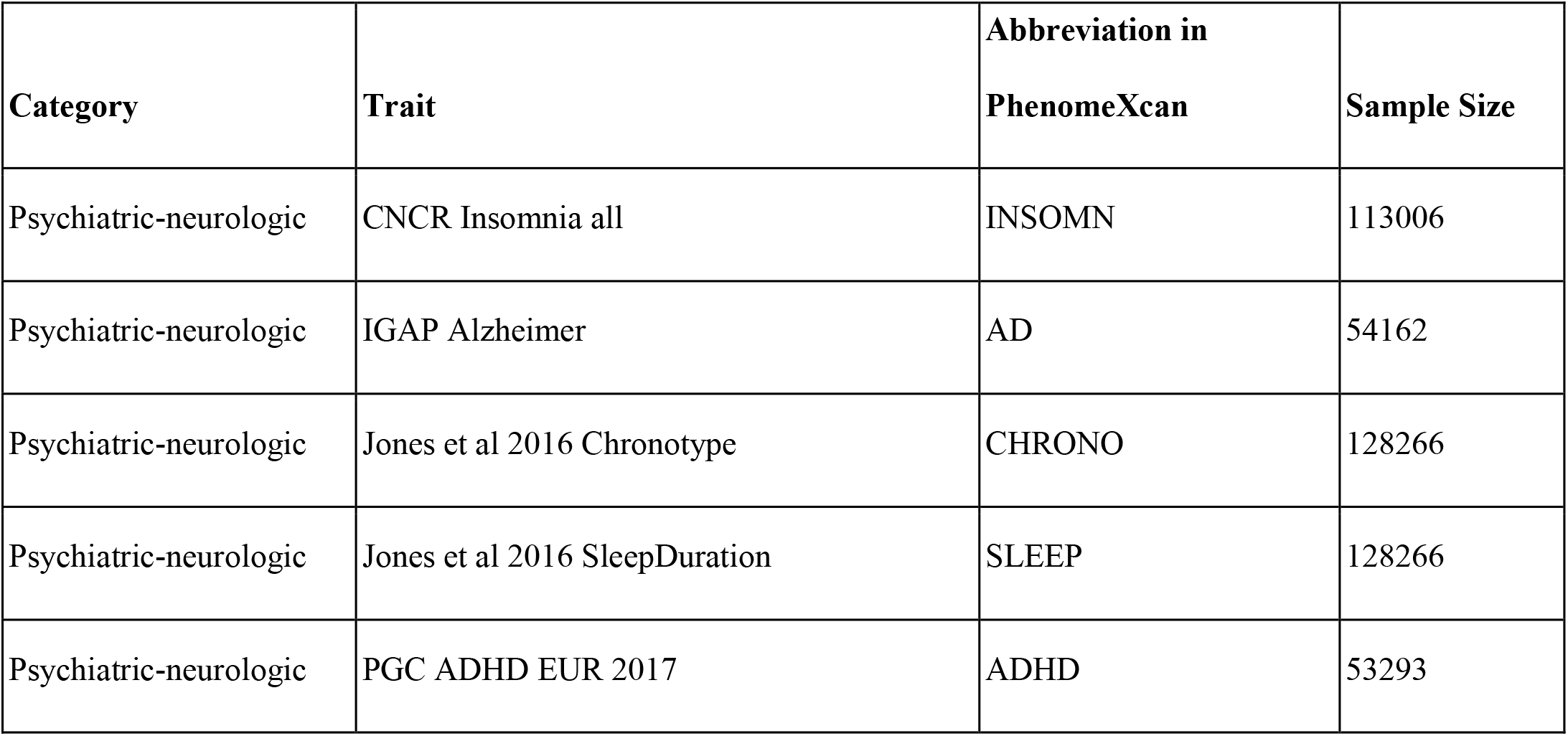
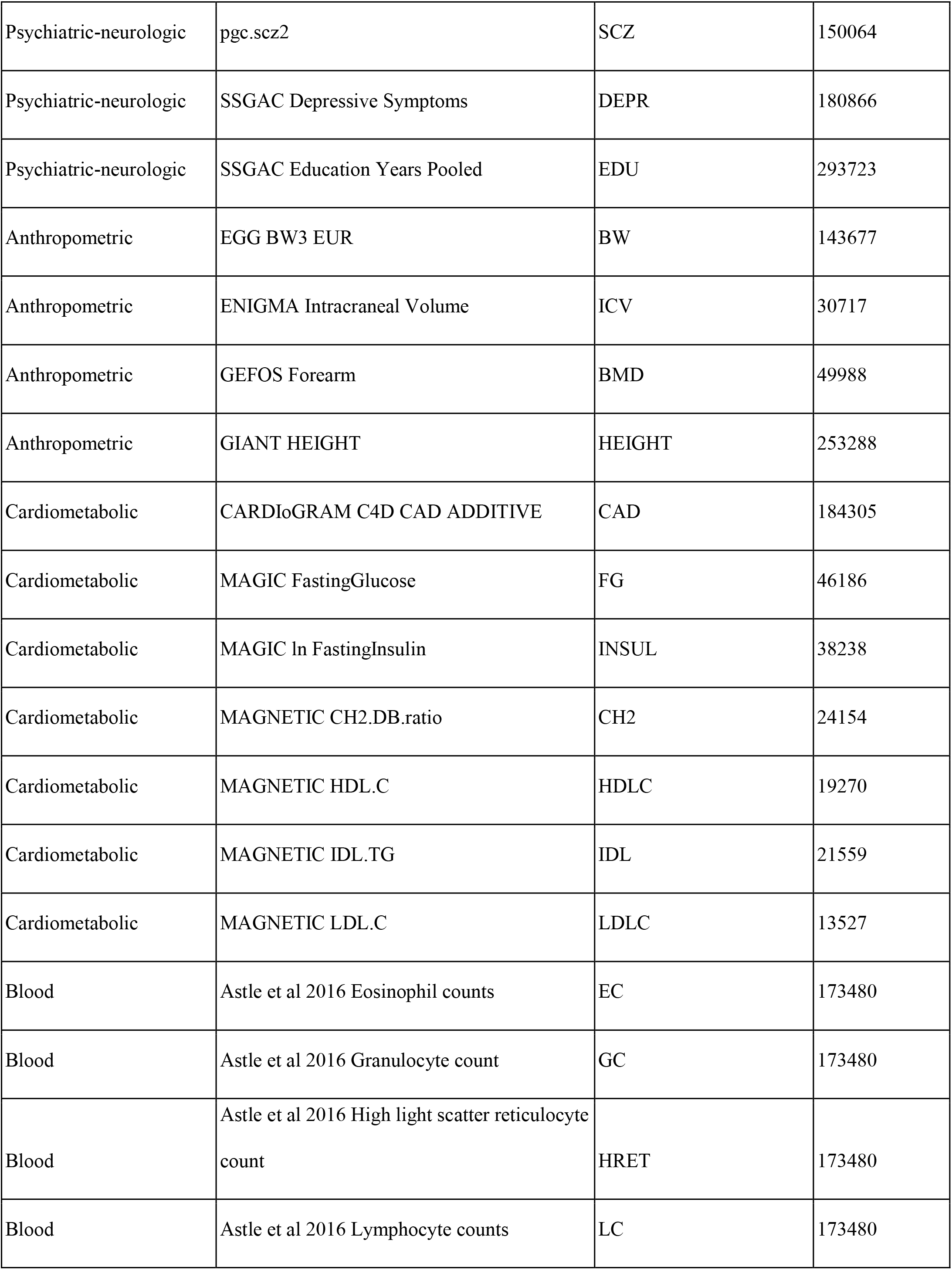
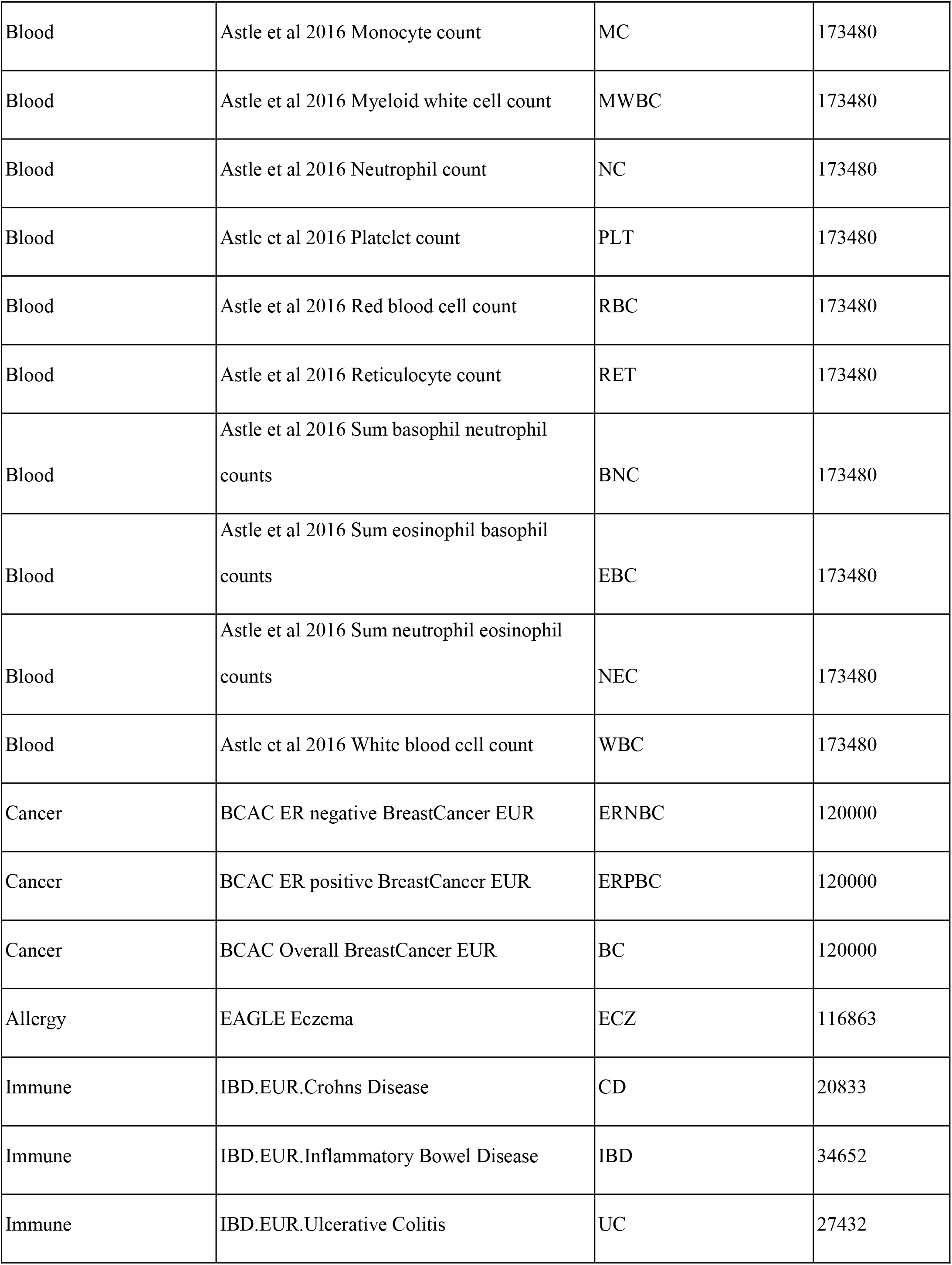
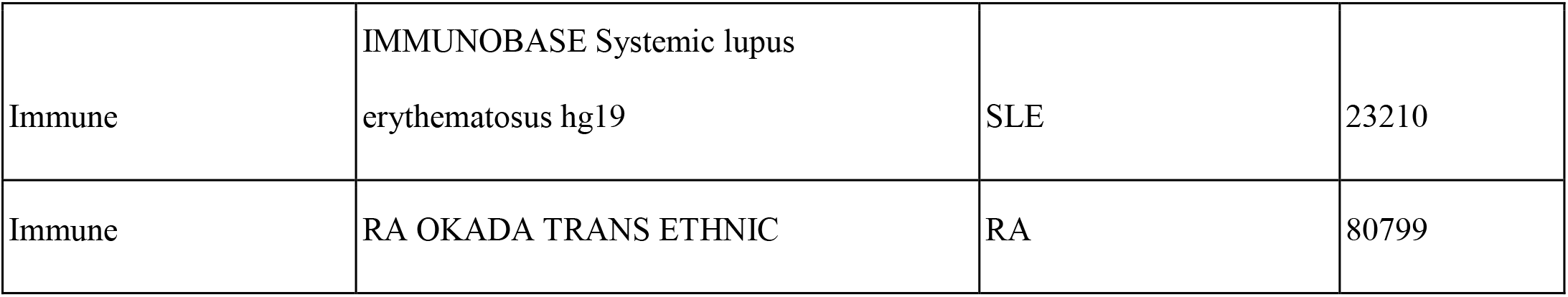
42 additional traits taken from GWAS studies for the development of PhenomeXcan. Traits are organized by trait category, data source, abbreviation in PhenomeXcan and number of subjects in the dataset.

**Supplementary Fig. S1:**
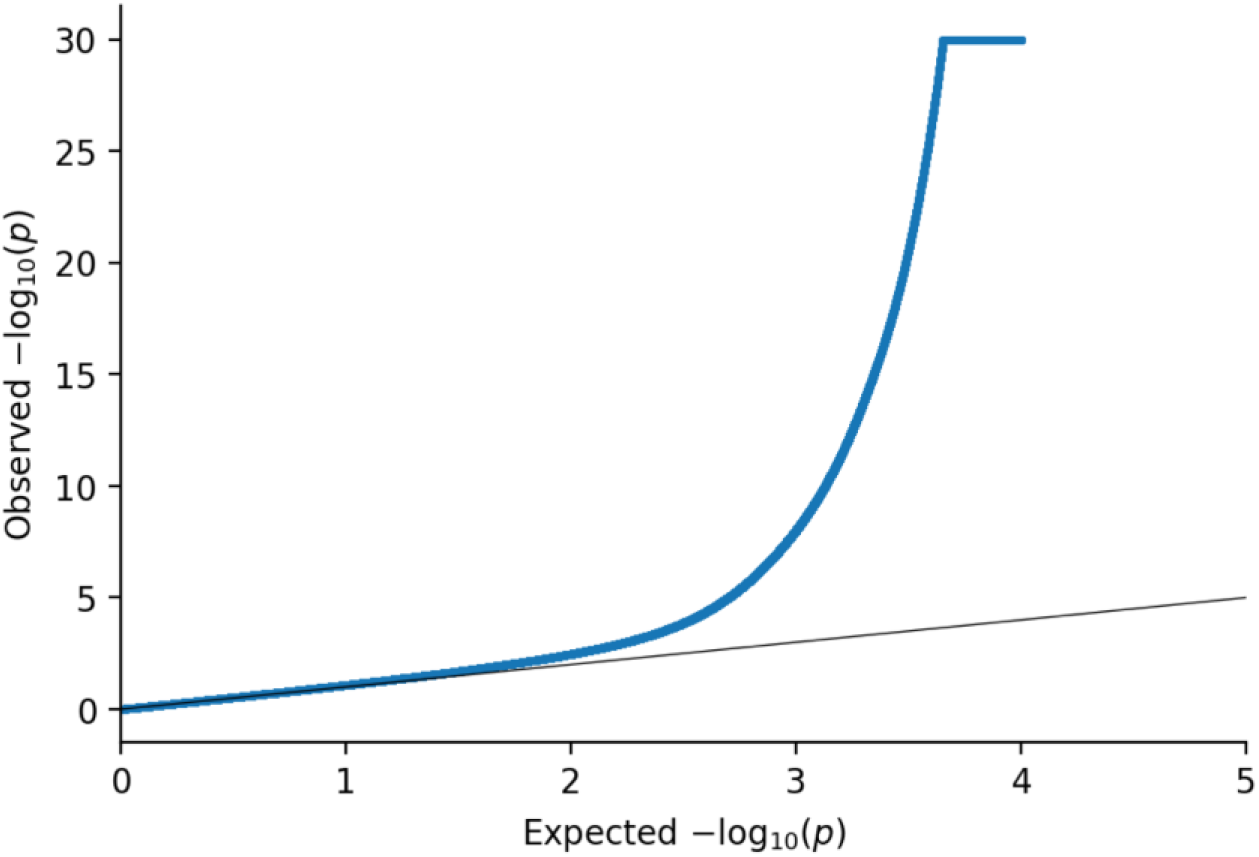
Quantile-quantile (QQ) plot of all associations in PhenomeXcan. The expected null distribution is plotted along the black diagonal, and the entire distribution of observed p-values is plotted in blue. We do not see evidence of systematic inflation given the initial consistency in expected and observed p-values. (To improve visualization, p-values are thresholded at −log10(p-value)=30.) The increase in the QQ plot for observed p-values can be seen with the extremely large number of associations tested.

**Supplementary Fig. S2:**
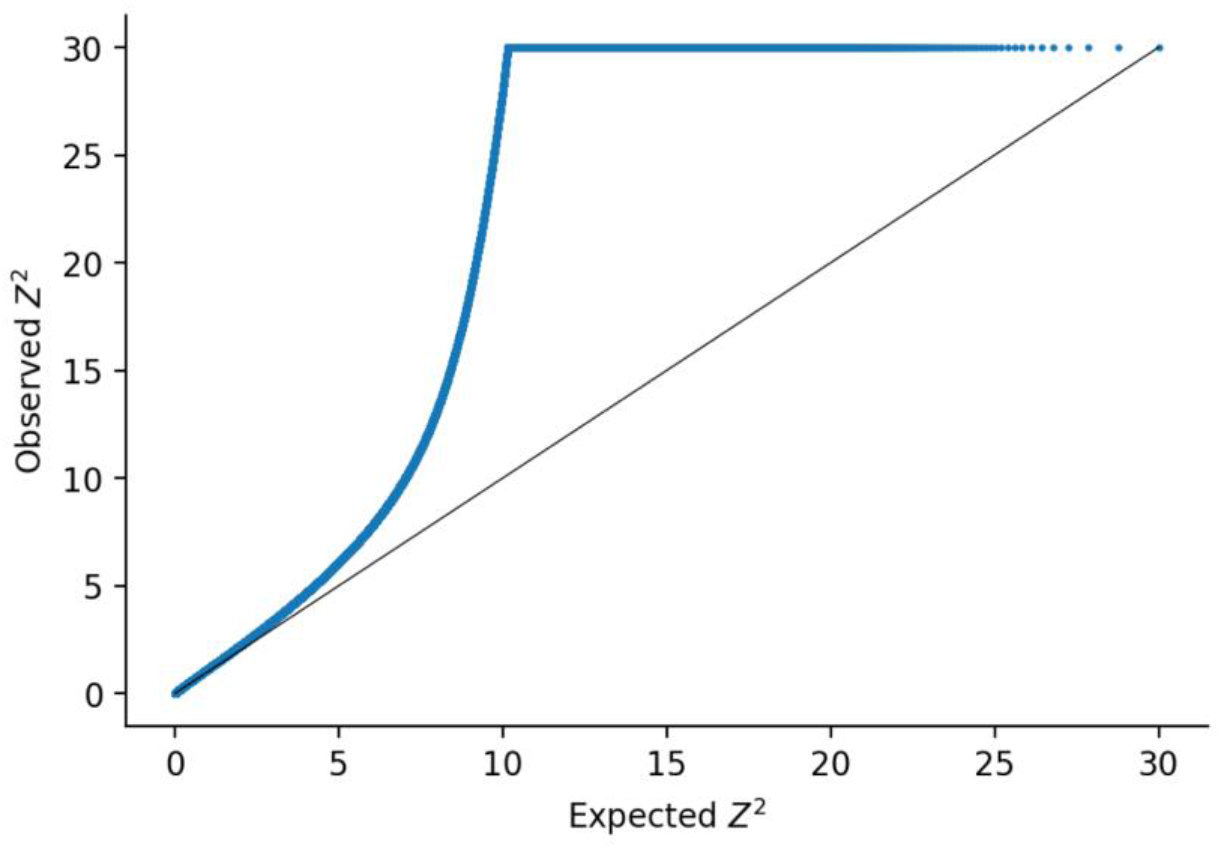
Quantile-quantile (QQ) plot of all associations in PhenomeXcan and ClinVar traits. The expected *χ*^2^null distribution is plotted along the black diagonal, and the entire distribution of observed *Z*^2^is plotted in blue. We do not see evidence of systematic inflation given the initial consistency in expected and observed p-values. (To improve visualization, *Z*^2^ are thresholded at 30.) The increase in the QQ plot for observed p-values can be seen with the extremely large number of associations tested (20.6 million) as well as the pleiotropy we identify with trait-trait associations in which multiple genes are involved. Z^2^ correspondence to percentiles were as follows: 95th percentile: Z^2^=4.45, 99th percentile: Z^2^=9.07, 99.9th percentile: Z^2^=214.45. A Z^2^ of 6 represents a Bonferroni-adjusted p-value of 0.05.

**Supplementary Fig. S3:**
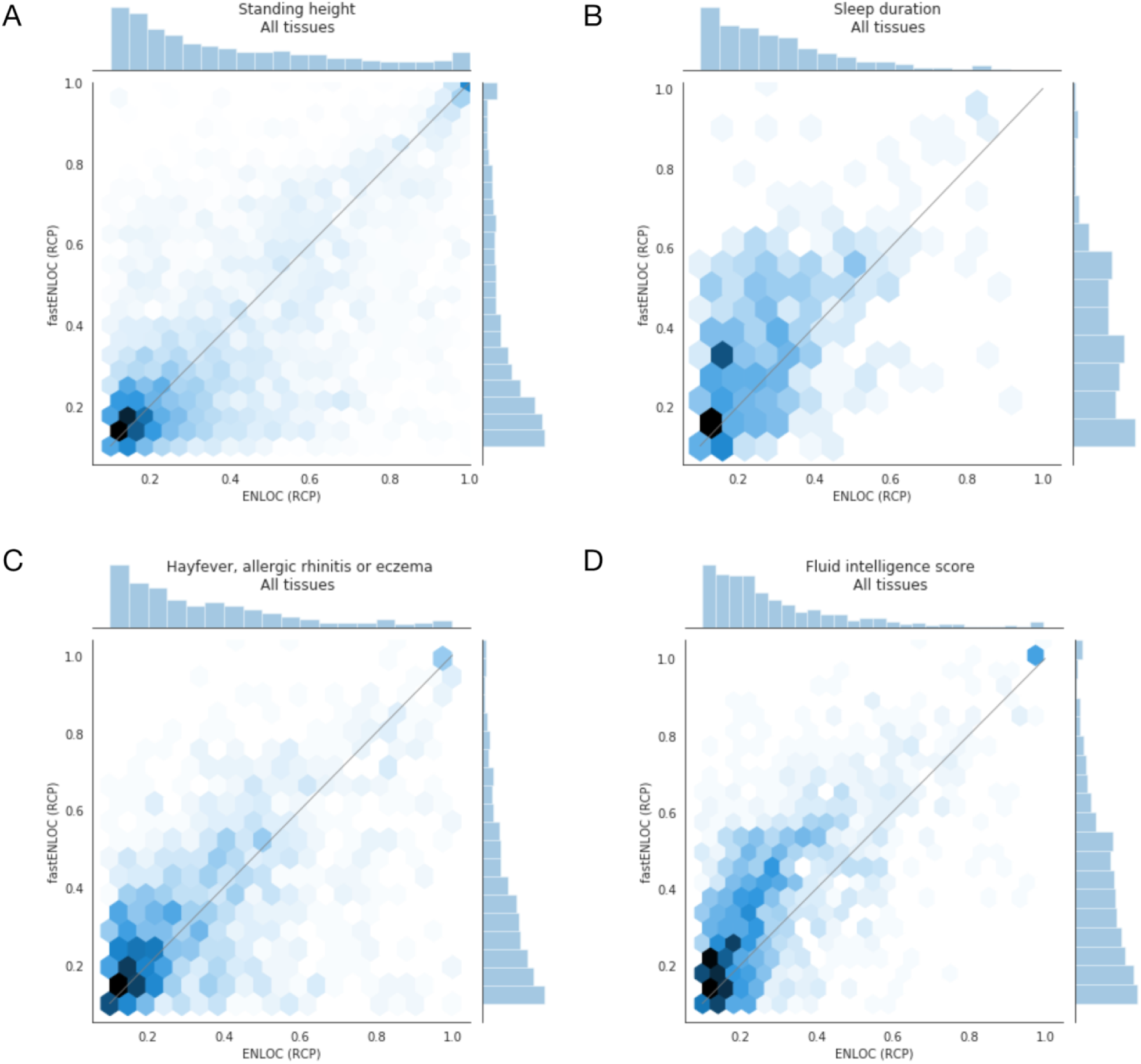
Joint histograms using hexagonal bins for the regional colocalization probability (RCP) agreement between fastENLOC and ENLOC. We analyzed the regional colocalization probabilities across traits between fastENLOC and ENLOC to assess their agreement. We found largely strong correlation between these methods, with the Spearman correlation coefficient for **(A) “** Standing height” = 0.61, **(B)** “Sleep duration” = 0.50 **, (C),** “Hayfever, allergic rhinitis or eczema” = 0.56 and **(D)** “Fluid intelligence score” = 0.65.

## List of GTEx Consortium Authors

Laboratory and Data Analysis Coordinating Center (LDACC): François Aguet^1^, Shankara Anand^1^, Kristin G Ardlie^1^, Stacey Gabriel^1^, Gad Getz^1,2^, Aaron Graubert^1^, Kane Hadley^1^, Robert E Handsaker^3,4,5^, Katherine H Huang^1^, Seva Kashin^3,4,5^, Xiao Li^1^, Daniel G MacArthur^4,6^, Samuel R Meier^1^, Jared L Nedzel^1^, Duyen Y Nguyen^1^, Ayellet V Segrè^1,7^, Ellen Todres^1^

Analysis Working Group (funded by GTEx project grants): François Aguet^1^, Shankara Anand^1^, Kristin G Ardlie^1^, Brunilda Balliu^8^, Alvaro N Barbeira^9^, Alexis Battle^10,11^, Rodrigo Bonazzola^9^, Andrew Brown^12,13^, Christopher D Brown^14^, Stephane E Castel^15,16^, Don Conrad^17,18^, Daniel J Cotter^19^, Nancy Cox^20^, Sayantan Das^21^, Olivia M de Goede^19^, Emmanouil T Dermitzakis^12,22,23^, Barbara E Engelhardt^24,25^, Eleazar Eskin^26^, Tiany Y Eulalio^27^, Nicole M Ferraro^27^, Elise Flynn^15, 16^, Laure Fresard^28^, Eric R Gamazon^20, 29, 30, 31^, Diego Garrido-Martín^32^, Nicole R Gay^19^, Gad Getz^1,2^, Aaron Graubert^1^, Roderic Guigó^32, 33^, Kane Hadley^1^, Andrew R Hamel^1, 7^, Robert E Handsaker^3,4,5^, Yuan He^10^, Paul J Homan^15^, Farhad Hormozdiari^1,34^, Lei Hou^1, 35^, Katherine H Huang^1^, Hae Kyung Im^9^, Brian Jo^24, 25^, Silva Kasela^15, 16^, Seva Kashin^3,4,5^, Manolis Kellis^1,35^, Sarah Kim-Hellmuth^15, 16, 36^, Alan Kwong^21^, Tuuli Lappalainen^15, 16^, Xiao Li^1^, Xin Li^28^, Yanyu Liang^9^, Daniel G MacArthur^4,6^, Serghei Mangul^26, 37^, Samuel R Meier^1^, Pejman Mohammadi^15, 16, 38, 39^, Stephen B Montgomery^19,28^, Manuel Muñoz-Aguirre^32, 40^, Daniel C Nachun^28^, Jared L Nedzel^1^, Duyen Y Nguyen^1^, Andrew B Nobel^41^, Meritxell Oliva^9,42^, YoSon Park^14,43^, Yongjin Park^1,35^, Princy Parsana^11^, Ferran Reverter^44^, John M Rouhana^1,7^, Chiara Sabatti^45^, Ashis Saha^11^, Ayellet V Segrè^1,7^, Andrew D Skol^9,46^, Matthew Stephens^47^, Barbara E Stranger^9,48^, Benjamin J Strober^10^, Nicole A Teran^28^, Ellen Todres^1^, Ana Viñuela^12,22,23,49^, Gao Wang^47^, Xiaoquan Wen^21^, Fred Wright^50^, Valentin Wucher^32^, Yuxin Zou^51^

Analysis Working Group (not funded by GTEx project grants): Pedro G Ferreira^52,53,54^, Gen Li^55^, Marta Melé^56^, Esti Yeger-Lotem^57,58^, Leidos Biomedical - Project Management: Mary E Barcus^59^, Debra Bradbury^60^, Tanya Krubit^60^, Jerey A McLean^60^, Liqun Qi^60^, Karna Robinson^60^ Nancy V Roche^60^, Anna M Smith^60^, Leslie Sobin^60^, David E Tabor^60^, Anita Undale^60^

Biospecimen collection source sites: Jason Bridge^61^, Lori E Brigham^62^, Barbara A Foster^63^, Bryan M Gillard^63^, Richard Hasz^64^, Marcus Hunter^65^, Christopher Johns^66^, Mark Johnson^67^, Ellen Karasik^63^, Gene Kopen^68^, William F Leinweber^68^, Alisa McDonald^68^, Michael T Moser^63^, Kevin Myer^65^, Kimberley D Ramsey^63^, Brian Roe^65^, Saboor Shad^68^, Jerey A Thomas^67,68^, Gary Walters^67^, Michael Washington^67^, Joseph Wheeler^66^

Biospecimen core resource: Scott D Jewell^69^, Daniel C Rohrer^69^, Dana R Valley^69^

Brain bank repository: David A Davis^70^, Deborah C Mash^70^

Pathology: Mary E Barcus^59^, Philip A Branton^71^, Leslie Sobin^60^

ELSI study: Laura K Barker^72^, Heather M Gardiner^72^, Maghboeba Mosavel^73^, Laura A Simino^72^

Genome Browser Data Integration & Visualization: Paul Flicek^74^, Maximilian Haeussler^75^, Thomas Juettemann^74^, W James Kent^75^, Christopher M Lee^75^, Conner C Powell^75^, Kate R Rosenbloom^75^, Magali Ru‑er^74^, Dan Sheppard^74^, Kieron Taylor^74^, Stephen J Trevanion^74^, Daniel R Zerbino^74^

eGTEx groups: Nathan S Abell^19^, Joshua Akey^76^, Lin Chen^42^, Kathryn Demanelis^42^, Jennifer A Doherty^77^, Andrew P Feinberg^78^, Kasper D Hansen^79^, Peter F Hickey^80^, Lei Hou^1,35^, Farzana Jasmine^42^, Lihua Jiang^19^, Rajinder Kaul^81,82^, Manolis Kellis^1,35^, Muhammad G Kibriya^42^, Jin Billy Li^19^, Qin Li^19^, Shin Lin^83^, Sandra E Linder^19^, Stephen B Montgomery^19,28^, Meritxell Oliva^9,42^, Yongjin Park^1,35^, Brandon L Pierce^42^, Lindsay F Rizzardi^84^, Andrew D Skol^9,46^, Kevin S Smith^28^, Michael Snyder^19^, John Stamatoyannopoulos^81,85^, Barbara E Stranger^9,48^, Hua Tang^19^, Meng Wang^19^

NIH program management: Philip A Branton^71^, Latarsha J Carithers^71,86^, Ping Guan^71^, Susan E Koester^87^, A. Roger Little^88^, Helen M Moore^71^, Concepcion R Nierras^89^, Abhi K Rao^71^, Jimmie B Vaught^71^, Simona Volpi^90^

Affiliations:

1. The Broad Institute of MIT and Harvard, Cambridge, MA, USA

2. Cancer Center and Department of Pathology, Massachusetts General Hospital, Boston, MA, USA

3. Department of Genetics, Harvard Medical School, Boston, MA, USA

4. Program in Medical and Population Genetics, The Broad Institute of Massachusetts Institute of Technology and Harvard University, Cambridge, MA, USA

5. Stanley Center for Psychiatric Research, Broad Institute, Cambridge, MA, USA

6. Analytic and Translational Genetics Unit, Massachusetts General Hospital, Boston, MA, USA

7. Ocular Genomics Institute, Massachusetts Eye and Ear, Harvard Medical School, Boston, MA, USA

8. Department of Biomathematics, University of California, Los Angeles, Los Angeles, CA, USA

9. Section of Genetic Medicine, Department of Medicine, The University of Chicago, Chicago, IL, USA

10. Department of Biomedical Engineering, Johns Hopkins University, Baltimore, MD, USA

11. Department of Computer Science, Johns Hopkins University, Baltimore, MD, USA

12. Department of Genetic Medicine and Development, University of Geneva Medical School, Geneva, Switzerland

13. Population Health and Genomics, University of Dundee, Dundee, Scotland, UK

14. Department of Genetics, University of Pennsylvania, Perelman School of Medicine, Philadelphia, PA, USA

15. New York Genome Center, New York, NY, USA

16. Department of Systems Biology, Columbia University, New York, NY, USA

17. Department of Genetics, Washington University School of Medicine, St. Louis, Missouri, USA

18. Department of Pathology & Immunology, Washington University School of Medicine, St. Louis, Missouri, USA

19. Department of Genetics, Stanford University, Stanford, CA, USA

20. Division of Genetic Medicine, Department of Medicine, Vanderbilt University Medical Center, Nashville, TN, USA

21. Department of Biostatistics, University of Michigan, Ann Arbor, MI, USA

22. Institute for Genetics and Genomics in Geneva (iGE3), University of Geneva, Geneva, Switzerland

23. Swiss Institute of Bioinformatics, Geneva, Switzerland

24. Department of Computer Science, Princeton University, Princeton, NJ, USA

25. Center for Statistics and Machine Learning, Princeton University, Princeton, NJ, USA

26. Department of Computer Science, University of California, Los Angeles, Los Angeles, CA, USA

27. Program in Biomedical Informatics, Stanford University School of Medicine, Stanford, CA, USA

28. Department of Pathology, Stanford University, Stanford, CA, USA

29. Data Science Institute, Vanderbilt University, Nashville, TN, USA

30. Clare Hall, University of Cambridge, Cambridge, UK

31. MRC Epidemiology Unit, University of Cambridge, Cambridge, UK

32. Centre for Genomic Regulation (CRG), The Barcelona Institute for Science and Technology, Barcelona, Catalonia, Spain

33. Universitat Pompeu Fabra (UPF), Barcelona, Catalonia, Spain

34. Department of Epidemiology, Harvard T.H. Chan School of Public Health, Boston, MA, USA

35. Computer Science and Artificial Intelligence Laboratory, Massachusetts Institute of Technology, Cambridge, MA, USA

36. Statistical Genetics, Max Planck Institute of Psychiatry, Munich, Germany

37. Department of Clinical Pharmacy, School of Pharmacy, University of Southern California, Los Angeles, CA, USA

38. Scripps Research Translational Institute, La Jolla, CA, USA

39. Department of Integrative Structural and Computational Biology, The Scripps Research Institute, La Jolla, CA, USA

40. Department of Statistics and Operations Research, Universitat Politècnica de Catalunya (UPC), Barcelona, Catalonia, Spain

41. Department of Statistics and Operations Research and Department of Biostatistics, University of North Carolina, Chapel Hill, NC, USA

42. Department of Public Health Sciences, The University of Chicago, Chicago, IL, USA

43. Department of Systems Pharmacology and Translational Therapeutics, University of Pennsylvania, Perelman School of Medicine, Philadelphia, PA, USA

44. Department of Genetics, Microbiology and Statistics, University of Barcelona, Barcelona. Spain.

45. Departments of Biomedical Data Science and Statistics, Stanford University, Stanford, CA, USA

46. Department of Pathology and Laboratory Medicine, Ann & Robert H. Lurie Children’s Hospital of Chicago, Chicago, IL, USA

47. Department of Human Genetics, University of Chicago, Chicago, IL, USA

48. Center for Genetic Medicine, Department of Pharmacology, Northwestern University, Feinberg School of Medicine, Chicago, IL, USA

49. Department of Twin Research and Genetic Epidemiology, King’s College London, London, UK

50. Bioinformatics Research Center and Departments of Statistics and Biological Sciences, North Carolina State University, Raleigh, NC, USA

51. Department of Statistics, University of Chicago, Chicago, IL, USA

52. Department of Computer Sciences, Faculty of Sciences, University of Porto, Porto, Portugal

53. Instituto de Investigação e Inovação em Sauúde, Universidade do Porto, Porto, Portugal

54. Institute of Molecular Pathology and Immunology, University of Porto, Porto, Portugal

55. Columbia University Mailman School of Public Health, New York, NY, USA

56. Life Sciences Department, Barcelona Supercomputing Center, Barcelona, Spain

57. Department of Clinical Biochemistry and Pharmacology, Ben-Gurion University of the Negev, Beer-Sheva, Israel

58. National Institute for Biotechnology in the Negev, Beer-Sheva, Israel

59. Leidos Biomedical, Frederick, MD, USA

60. Leidos Biomedical, Rockville, MD, USA

61. UNYTS, Bualo, NY, USA

62. Washington Regional Transplant Community, Annandale, VA, USA

63. Therapeutics, Roswell Park Comprehensive Cancer Center, Bualo, NY, USA

64. Gift of Life Donor Program, Philadelphia, PA, USA

65. LifeGift, Houston, TX, USA

66. Center for Organ Recovery and Education, Pittsburgh, PA, USA

67. LifeNet Health, Virginia Beach, VA. USA

68. National Disease Research Interchange, Philadelphia, PA, USA

69. Van Andel Research Institute, Grand Rapids, MI, USA

70. Department of Neurology, University of Miami Miller School of Medicine, Miami, FL, USA

71. Biorepositories and Biospecimen Research Branch, Division of Cancer Treatment and Diagnosis, National Cancer Institute, Bethesda, MD, USA

72. Temple University, Philadelphia, PA, USA

73. Virgina Commonwealth University, Richmond, VA, USA

74. European Molecular Biology Laboratory, European Bioinformatics Institute, Hinxton, United Kingdom v

75. Genomics Institute, UC Santa Cruz, Santa Cruz, CA, USA

76. Carl Icahn Laboratory, Princeton University, Princeton, NJ, USA

77. Department of Population Health Sciences, The University of Utah, Salt Lake City, Utah, USA

78. Schools of Medicine, Engineering, and Public Health, Johns Hopkins University, Baltimore, MD, USA

79. Department of Biostatistics, Bloomberg School of Public Health, Johns Hopkins University, Baltimore, MD, USA

80. Department of Medical Biology, The Walter and Eliza Hall Institute of Medical Research, Parkville, Victoria, Australia

81. Altius Institute for Biomedical Sciences, Seattle, WA, USA

82. Division of Genetics, University of Washington, Seattle, WA, University of Washington, Seattle, WA, USA

83. Department of Cardiology, University of Washington, Seattle, WA, USA

84. HudsonAlpha Institute for Biotechnology, Huntsville, AL, USA

85. Genome Sciences, University of Washington, Seattle, WA, USA

86. National Institute of Dental and Craniofacial Research, Bethesda, MD, USA

87. Division of Neuroscience and Basic Behavioral Science, National Institute of Mental Health, National Institutes of Health, Bethesda, MD, USA

88. National Institute on Drug Abuse, Bethesda, MD, USA

89. Office of Strategic Coordination, Division of Program Coordination, Planning and Strategic Initiatives, O-ce of the Director, National Institutes of Health, Rockville, MD, USA

90. Division of Genomic Medicine, National Human Genome Research Institute, Bethesda, MD, USA

This work was funded by GTEx program grants: HHSN268201000029C (F.A., K.G.A., A.V.S., X.Li., E.T., S.G., A.G., S.A., K.H.H., D.Y.N., K.H., S.R.M., J.L.N.), 5U41HG009494 (F.A., K.G.A.), 10XS170 (Subcontract to Leidos Biomedical) (W.F.L., J.A.T., G.K., A.M., S.S., R.H., G.Wa., M.J., M.Wa., L.E.B., C.J., J.W., B.R., M.Hu., K.M., L.A.S., H.M.G., M.Mo., L.K.B.), 10XS171 (Subcontract to Leidos Biomedical) (B.A.F., M.T.M., E.K., B.M.G., K.D.R., J.B.), 10ST1035 (Subcontract to Leidos Biomedical) (S.D.J., D.C.R., D.R.V.), R01DA006227-17 (D.C.M., D.A.D.), Supplement to University of Miami grant DA006227. (D.C.M., D.A.D.), HHSN261200800001E (A.M.S., D.E.T., N.V.R., J.A.M., L.S., M.E.B., L.Q., T.K., D.B., K.R., A.U.), R01MH101814 (M.M-A., V.W., S.B.M., R.G., E.T.D., D.G-M., A.V.), U01HG007593 (S.B.M.), R01MH101822 (C.D.B.), U01HG007598 (M.O., B.E.S.), as well as other funding sources: R01MH106842 (T.L., P.M., E.F., P.J.H.), R01HL142028 (T.L., Si.Ka., P.J.H.), R01GM122924 (T.L., S.E.C.), R01MH107666 (H.K.I.), P30DK020595 (H.K.I.), UM1HG008901 (T.L.), R01GM124486 (T.L.), R01HG010067 (Y.Pa.), R01HG002585 (G.Wa., M.St.), Gordon and Betty Moore Foundation GBMF 4559 (G.Wa., M.St.), 1K99HG009916-01 (S.E.C.), R01HG006855 (Se.Ka., R.E.H.), BIO2015-70777-P, Ministerio de Economia y Competitividad and FEDER funds (M.M-A., V.W., R.G., D.G-M.), NIH CTSA grant UL1TR002550-01 (P.M.), Marie-Skłodowska Curie fellowship H2020 Grant 706636 (S.K-H.), R35HG010718 (E.R.G.), FPU15/03635, Ministerio de Educación, Cultura y Deporte (M.M-A.), R01MH109905, 1R01HG010480 (A.Ba.), Searle Scholar Program (A.Ba.), R01HG008150 (S.B.M.), 5T32HG000044-22, NHGRI Institutional Training Grant in Genome Science (N.R.G.), EU IMI program (UE7-DIRECT-115317-1) (E.T.D., A.V.), FNS funded project RNA1 (31003A_149984) (E.T.D., A.V.), DK110919 (F.H.), F32HG009987 (F.H.)

F.A. is an inventor on a patent application related to TensorQTL; S.E.C. is a co-founder, chief technology officer and stock owner at Variant Bio; E.R.G. is on the Editorial Board of Circulation Research, and does consulting for the City of Hope / Beckman Research Institut; E.T.D. is chairman and member of the board of Hybridstat LTD.; B.E.E. is on the scientific advisory boards of Celsius Therapeutics and Freenome; G.G. receives research funds from IBM and Pharmacyclics, and is an inventor on patent applications related to MuTect, ABSOLUTE, MutSig, POLYSOLVER and TensorQTL; S.B.M. is on the scientific advisory board of Prime Genomics Inc.; D.G.M. is a co-founder with equity in Goldfinch Bio, and has received research support from AbbVie, Astellas, Biogen, BioMarin, Eisai, Merck, Pfizer, and Sanofi-Genzyme; H.K.I. has received speaker honoraria from GSK and AbbVie.; T.L. is a scientific advisory board member of Variant Bio with equity and Goldfinch Bio. P.F. is member of the scientific advisory boards of Fabric Genomics, Inc., and Eagle Genomes, Ltd. P.G.F. is a partner of Bioinf2Bio.

## Notes

#### Summary of Updates

Text has been revised.

https://github.com/hakyimlab/phenomexcan

http://phenomexcan.org/

